# A zinc knuckle gene on the Y chromosome (*zkY*) determines sex in Atlantic cod (*Gadus morhua*)

**DOI:** 10.1101/362376

**Authors:** Tina Graceline Kirubakaran, Øivind Andersen, Maria Cristina De Rosa, Terese Andersstuen, Kristina Hallan, Matthew Peter Kent, Sigbjørn Lien

## Abstract

The genetic mechanisms determining sex in teleost fishes are highly variable, ranging from a single gene to complex patterns of polygenic regulation. The master sex determining gene has only been identified in very few species and there is no information about the gene in the superorder Paracanthopterygii that includes the codfishes, toadfishes and anglerfishes. Here we characterize a male-specific region of 9 kb on linkage group 11 in Atlantic cod (*Gadus morhua*) harboring a single gene named *zkY* for zinc knuckle on the Y chromosome. A diagnostic PCR test of phenotypically sexed males and females of Atlantic cod confirmed the sex-specific nature of the Y-sequence. We searched for autosomal gene copies of *zkY* and identified twelve highly similar genes, of which eight (*zk1-zk8*) code for proteins containing the zinc knuckle motif. 3D structure modelling suggests that the amino acid changes observed in six of the eight copies might influence the putative RNA-binding specificity. Cod zkY and the autosomal proteins zk1 and zk2 possess an identical zinc knuckle structure, but only the Y-specific gene *zkY* was expressed at high levels in the developing larvae before the onset of sex differentiation. We propose that cod *zkY* functions as the master masculinization gene by coding for a suppressor of germ cell division in males. PCR amplification of Y-sequences in Arctic cod (*Arctogadus glacialis*) and Greenland cod (*Gadus macrocephalus ogac*) suggests that this novel sex determining mechanism emerged in codfishes more than 7.5 million years ago.

**Author Summary:** Studying the diverse sex determining genes in teleost fish may contribute to increase our understanding of sex chromosome evolution in vertebrates. To date, no sex determinant is known for the superorder Paracanthopterygii, comprising about 1340 species, including the commercially important Atlantic cod. In this study we characterize a Y-specific region of 9 kb on linkage group 11 containing a single gene named *zkY* for zinc knuckle on the Y chromosome. The gene is transcribed at high levels in larvae before commencement of sex differentiation and encodes a novel zinc knuckle protein that putatively binds RNA target sequences. We propose that cod zkY suppresses germ cell proliferation in the developing males by interacting with the germ-cell specific RNA regulatory network.

## Introduction

The origin and evolution of sex chromosomes from autosomes, and the mechanism of sex determination have long been subjects of interest to biologists. However, the highly dimorphic sex chromosomes in eutherian mammalian and birds are degenerated with extensive gene losses [1-3]. In teleost fishes, cytogenetically different sex chromosomes are found in less than 10% of the species examined [4, 5], and their recent origin in several lineages makes them good models to study the early stages of divergence. The frequent turnover of sex chromosomes seems to be associated with the variety of sex determining genes, even in closely related species [4-6]. For example, in medaka fishes (genus *Oryzias*), the Y-chromosomal *dmY* or *dmrt1bY* copy determines the sex in *O. latipes* and *O. curvinotus*, while testicular differentiation in *O. dancena* and in *O. luzonensis* is initiated by male-specific regulatory elements upstream of *sox3* and *gsdf* (gonadal soma-derived factor), respectively [7-10]. The various mechanisms of genetic sex determination in fish range from sex–specific alleles of the anti-Müllerian hormone (Amh) in Nile tilapia (*Oreochromis niloticus*) and the Amh type 2 receptor (*Amhr2*) in tiger pufferfish (*Takifugu rubripes*) [11, 12] to complex polygenic regulation involving several genomic regions as shown in the cichlid *Astatotilapia burtoni* [13]. The diversity of mechanisms is further increased by the insertion of X- and Y-sequences modulating the expression of the neighboring master sex determinant gene, as found in the sablefish (*Anoplopoma fimbria*) [14].

Sexual differentiation in vertebrates is initially marked by highly increased proliferation of the germ cells in the presumptive ovaries compared to that in testes [6, 15-17]. The time course of the sex-dimorphic germ cell proliferation varies greatly among teleost species and occurs around hatching in medaka [18], at the start-feeding stage in Atlantic cod [19] and in late juvenile stages in sea bass (*Dicentrarchus labrax*) [20]. Male germ cell divisions are inhibited by Dmy in the medaka *O. latipes* [21], and the sex determinants Gdsf, Amh and Amhr2 of the TGFβ signaling pathway might play similar roles in inducing sex differentiation in other species [6]. However, the requirement of germ cells for gonadal development appears to be vary among teleost species [22-24] reflecting that sex determination might be triggered by alternative mechanisms. Most sex determinants found in vertebrates are thought to have acquired this role by being recruited from conserved downstream regulators of gonadal differentiation, although the function of some actors differs among lineages [6, 25-27]. An exception is the novel salmonid sdY; a truncated male-specific copy of the interferon regulatory factor 9 (irf9), which is an immune-related gene not associated with sex [28]. Knowledge about how genes are functionalized and incorporated at the top of the sex regulatory cascade should broaden our understanding of the genetic regulation of sex determination and elucidate whether there are constraints on the types of genes that can be co-opted as master control switches.

Atlantic cod is an economically important cold-water marine species widely distributed in the North Atlantic Ocean. Due to the decline in many cod stocks, cod farming has attracted interest but production is hampered by precocious early sexual maturation, particularly in males. This could partly be solved by production of all-female triploid stocks [4, 29, 30]. Gynogenetic and sex-reversed cod populations have demonstrated a XX-XY sex determination system [19, 31, 32], but karyotyping of Atlantic cod and the closely related European hake (*Merluccius merluccius*) failed to reveal sex-linked chromosome heteromorphism [33, 34]. Recently, whole genome sequence data from wild Atlantic cod was used to identify genotypes segregating closely with a XX-XY system in six putative sex determining regions distributed across five linkage groups [35]. Here we use whole genome re-sequencing data from 49 males and 53 females, together with long-read sequence data and Sanger sequencing of targeted PCR products, to characterize a Y-sequence of 9,149 base pairs on LG11 harboring a single gene named zinc knuckle on the Y chromosome gene *(zkY)*. Gene expression data from early development stages and modeling of the zinc knuckle structure, support the identification of the Y-specific *zkY* as the likely major masculinization gene in Atlantic cod.

## Results and Discussion

### Identification of the sex determining region

Two independent approaches were used to identify the sex determining region in Atlantic cod. Firstly, we searched whole genome sequence data from males and females for SNPs that segregate according to an XX-XY system. Illumina short-reads (approximately 10X coverage per individual) generated from 49 males and 53 females were mapped to the in-house developed gadMor2.1 genome assembly (S1 File), followed by variant detection. When applying stringent criteria demanding that gender specific variants should be heterozygous in all 49 males and homozygous in all 53 females, we detected 9 variants all distributed within a 15 kb region on LG11 (S2 File and Fig 1b). BLAST alignments revealed that these variants fell inside the 55 kb region previously reported as a likely candidate sex determining locus [35]. Similar results were obtained when applying this mapping approach against the gadMor2.0 assembly [36] leading us to conclude that the sex determining locus in Atlantic cod maps to this region on LG11. Secondly, we analyzed resequencing data to identify positions in the genome that displayed significant differences in read depth between males and females. Two regions showed an almost complete absence of female reads while displaying >500 reads from males, i.e. Y-specific characteristics. These regions were separated by an intervening sequence displaying roughly 2-times coverage in females versus males, i.e. a possible X-sequence. Both the Y- and X-sequence regions fell within an interval flanked by the nine sex-linked markers on LG11 lending further support to the significance of this region in sex determination (Fig 1a). Closer examination of the architecture of this region using IGV [37] revealed that despite a high read-depth, there was no evidence of single reads bridging, or read-pairs spanning, these differential read-depth sub-regions. Collectively these anomalies are suggestive of assembly errors in the initial gadMor2.1 assembly as a hybridization of X- and Y-sequences.

**Fig 1.**
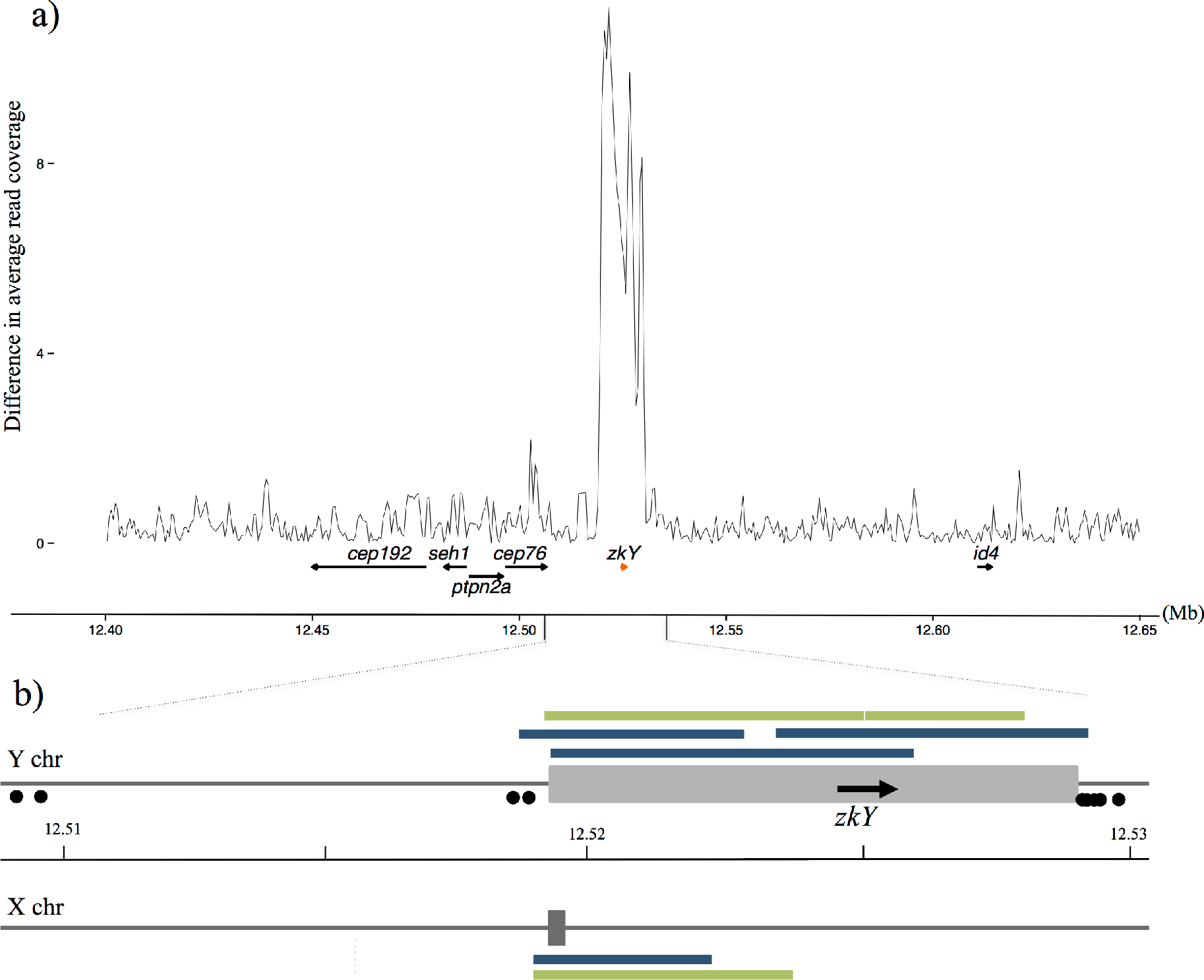
Genomic organization of the Y- and X-sequences on LG11 in Atlantic cod. (a) Comparison of read depth between male (n=49) and female (n=53) samples reveals an excess of Y-specific reads across a 9 kb interval on LG11 in the gadMor2.1 assembly. Positions of cod *zkY* and the neighboring genes are indicated. (b) The Y- and X-sequences (grey boxes), including the Y-specific *zkY* gene, and the nine polymorphisms heterozygous in all males and homozygous in all females (black dots). Long reads confirming assembly integrity were generated using Oxford Nanopore (green) and Pacific Biosciences (blue) sequencing technologies.

In order to resolve this complex region we performed nested, long-range PCR to generate a Y-specific fragment, which was then sequenced using an Illumina MiSeq. Assembly construction using Gap Filler generated a 6kb sequence, which was integrated into the gadMor2.1 assembly to produce a complete Y-sequence of 9,149bp (S3 File). The sequencing revealed an imperfect repeated element positioned close to the end of the Y-sequence (see underlined sequence in S3 File). An X-sequence of 425bp was constructed by Sanger sequencing PCR fragments generated from females using primers flanking the Y-sequence. To confirm the integrity of these manually curated X- and Y-sequences we aligned publicly available long-read data (Pacific BioSciences) derived from the male used to generate the reference (accession numbers at ENA (http://www.ebi.ac.uk/ena), ERX1787826-ERX1787972) and long reads generated in-house using an Oxford Nanopore MinION device, also from a male. Although overall coverage was low, we were able to identify long-reads that aligned specifically to either the Y- or X-sequences and, in combination, spanned their entire lengths (S4 File and Fig. 1b).

### The Y-sequence contains a single gene

X- and Y-sequences of the gadMor2.1 assembly were annotated using standard gene-model predictive software and RNA-Seq data produced from early larval stages together with publicly available RNA-Seq data from Atlantic cod (See Material and Methods). This revealed a single gene named *zkY* in the Y-sequence (S3 File and Fig 1), which is inserted in a synteny block being highly conserved in teleosts and in spotted gar (*Lepisosteus oculatus*). The intronless cod *zkY* codes for a novel zinc knuckle protein characterized by the zinc knuckle consensus sequence Cys-X2-Cys-X4-His-X4-Cys (X= any amino acid). This motif is mainly found in the RNA-binding retroviral nucleocapsid (NC) proteins, but also in various eukaryotic proteins binding single-stranded nucleotide targets [38-41]. The flanking basic residues bind nucleic acids non-specifically through electrostatic interactions with the phosphodiester backbone of nucleic acids [42].

Zinc knuckle proteins are members of the large family of zinc finger proteins possessing a versatility of tetrahedral Cys- and His-containing motifs that bind to DNA and RNA target sites [43, 44]. The DNA-binding domain of the DMRT transcription factors, including the sex determinant Dmy in medaka, consists of intertwined CCHC and HCCC motifs binding in the minor groove of DNA [45]. The zinc finger protein ZFAND3 is essential for spermatogenesis in mice and the polymorphic tilapia *zfand3* was recently mapped in the sex determining locus in tilapia [46, 47]. Hence, the recruitment of a zinc finger, or zink knuckle, proteins as the sex determinant seems to have occurred several times in teleosts.

### Multiple autosomal copies of cod zkY

The sex determining gene in several teleosts exhibits an autosomal copy that differ from the male-specific gene in spatio-temporal expression patterns and/or functionality [26, 48-51]. We identified 12 autosomal copies of cod *zkY* mapping to seven different linkage groups and one unassembled scaffold. Premature stop codons and single base indels were found in four of the genes encoding putatively non-functional proteins lacking the zinc knuckle domain. The remaining eight genes code for zinc knuckle proteins named zk1 to zk8, which differ in length from 143 to 633 amino acids (S5 Table). Sequence alignment revealed several amino acid substitutions in the zinc knuckle domain in addition to the variable size of an imperfect repeat of basic residues preceding the domain (S6 File and Fig 2A).

**Fig 2.**
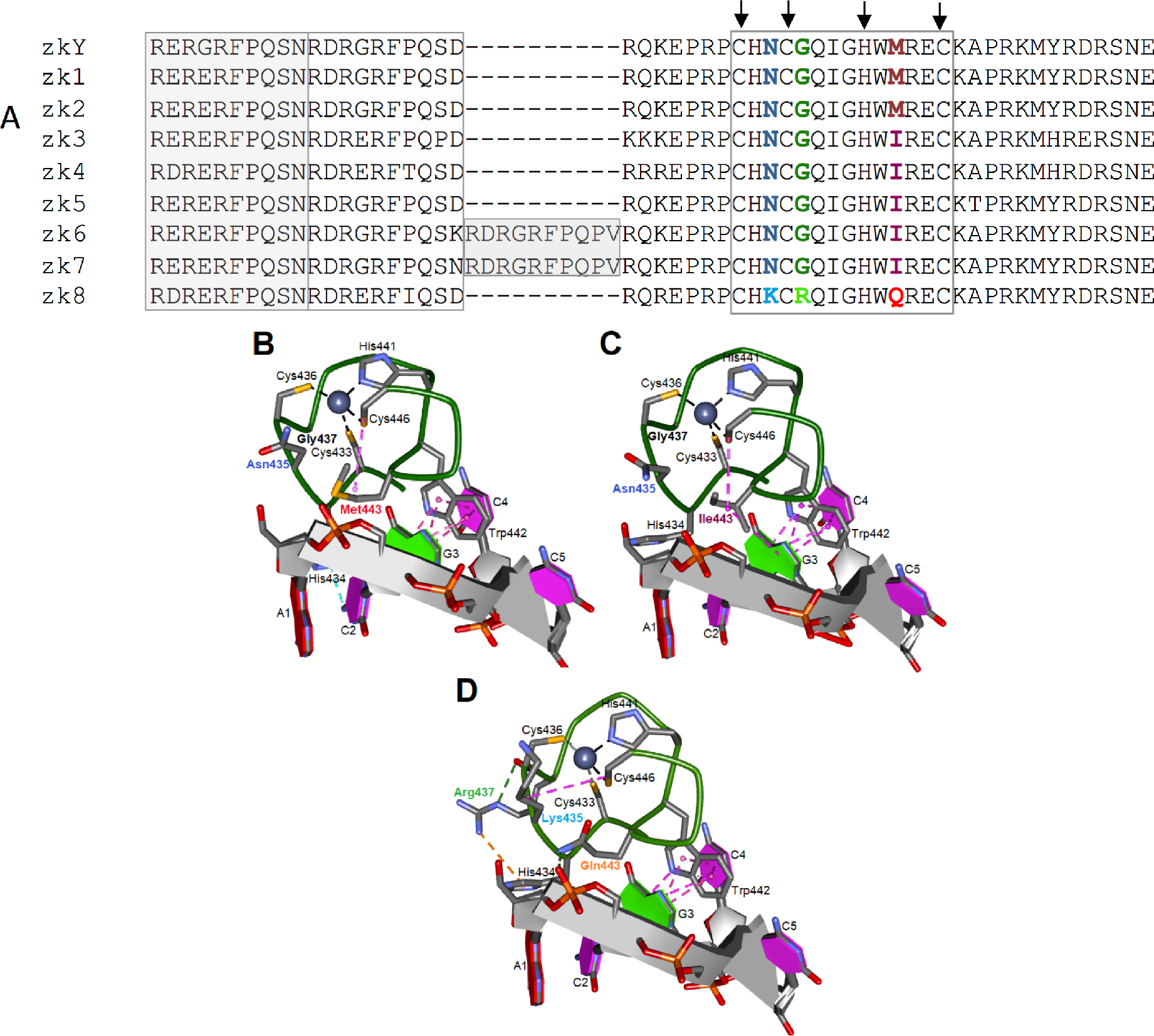
Amino acid substitutions in the zinc knuckle domain in cod zkY and its autosomal copies. A) Sequence alignment of the zinc knuckle domain (boxed) and flanking regions in cod zkY and the eight autosomal protein variants. The characteristic Cys-Cys-His-Cys residues of the zinc knuckle are arrowed, substituted amino acids are indicated by colors and correspond to the labelled positions 435, 437, and 443 in B-D. Repeated segments adjacent to the zinc knuckle are shaded. B-D. Modeled structure of the three different zinc knuckle domains of cod zkY and the autosomal proteins. The Zn^2+^ ion is displayed together with the oligonucleotide d(ACGCC) template (see Methods). B) Met443 variant of zkY, zk1, zk2, C) Ile443 variant of zk3-zk7, D) Lys435-Arg437-Gln443 variant of zk8. Hydrogen bonds are indicated by dotted green lines, Pi-donor hydrogen bonds in light blue, hydrophobic interactions by dotted magenta lines, and electrostatic bonds by dotted orange lines.

Functional implications of the amino acid substitutions in the zinc knuckle domain of the autosomal proteins was inferred by exploring the predicted 3D structure and the interactions with a putative RNA oligonucleotide target. The suitability of the d(ACGCC) sequence in the structure modeling was supported by the stacking of the conserved Trp442 between the two bases G3 and C4 in agreement with the interactions between specific base moieties and Trp at the corresponding position in NC proteins [52-54]. The three cysteins Cys443, Cys436 and Cys446 together with His441 coordinate the tetrahedral binding of the Zn^2+^ ion in the modelled cod zinc knuckles. While the replacement of Met433 in zkY, zk1 and zk2 with the Ile residue in the four zk3-zk7 proteins maintains the hydrophobic interactions with Cys446, the Ile433 residue is predicted to interact with the G3 base of the pentanucleotide (Fig 2C). The RNA-binding specificity of human HIV-1 was consistently altered by the Met->Val and Met->Lys substitutions in the same position of the second zinc knuckle [55]. The Asn435Lys change in the zk8 protein may affect the tetrahedral Zn^2+^ ion coordination by interacting with Cys446, while Gln443 is predicted to form a hydrogen bond with the phosphate group connecting A1 and C2 (Fig 2D). Additionally, electrostatic interactions are predicted between Arg437 and His434, which are moved away from its C2 contacts observed in the other variants.

Altogether, the predicted alterations in the contacts between the replaced amino acids and the interactions with the oligonucleotide sequence template suggest functional differences in the putative RNA-binding activity of the zinc knuckle proteins examined. In addition, the functional role played by the flanking basic residues in the non-specific binding of nucleic acids [43] suggests that these interactions might be altered by the extended flanking repeat in the zk6 and zk7 proteins. Based on the identical zinc knuckle domain and the similar flanking sequences in the three proteins zkY, zk1 and zk2, we decided to compare the larval expression of these coding genes.

### Increased transcription of cod *zkY* prior to onset of sex differentiation

A prerequisite for being a sex determining gene is to show pronounced transcription levels prior to onset of sex differentiation, which in Atlantic cod likely occurs at start feeding around 35 days post hatching (dph) based on the high number of germ cells in females compared to males together with the female-biased expression of *cyp19a1a* [19, 56]. RNA-Seq analysis of cod *zkY* and the autosomal *zk1* and *zk2* copies showed that the three genes were expressed at variable levels from hatching until 22 dph at which stage the transcription levels of *zkY* increased and stabilized at relatively high levels from 30 dph onwards (Fig 3). In contrast, the larval expression of *zk1* and *zk2* was almost undetectable or very low from 11 dph.

**Fig 3.**
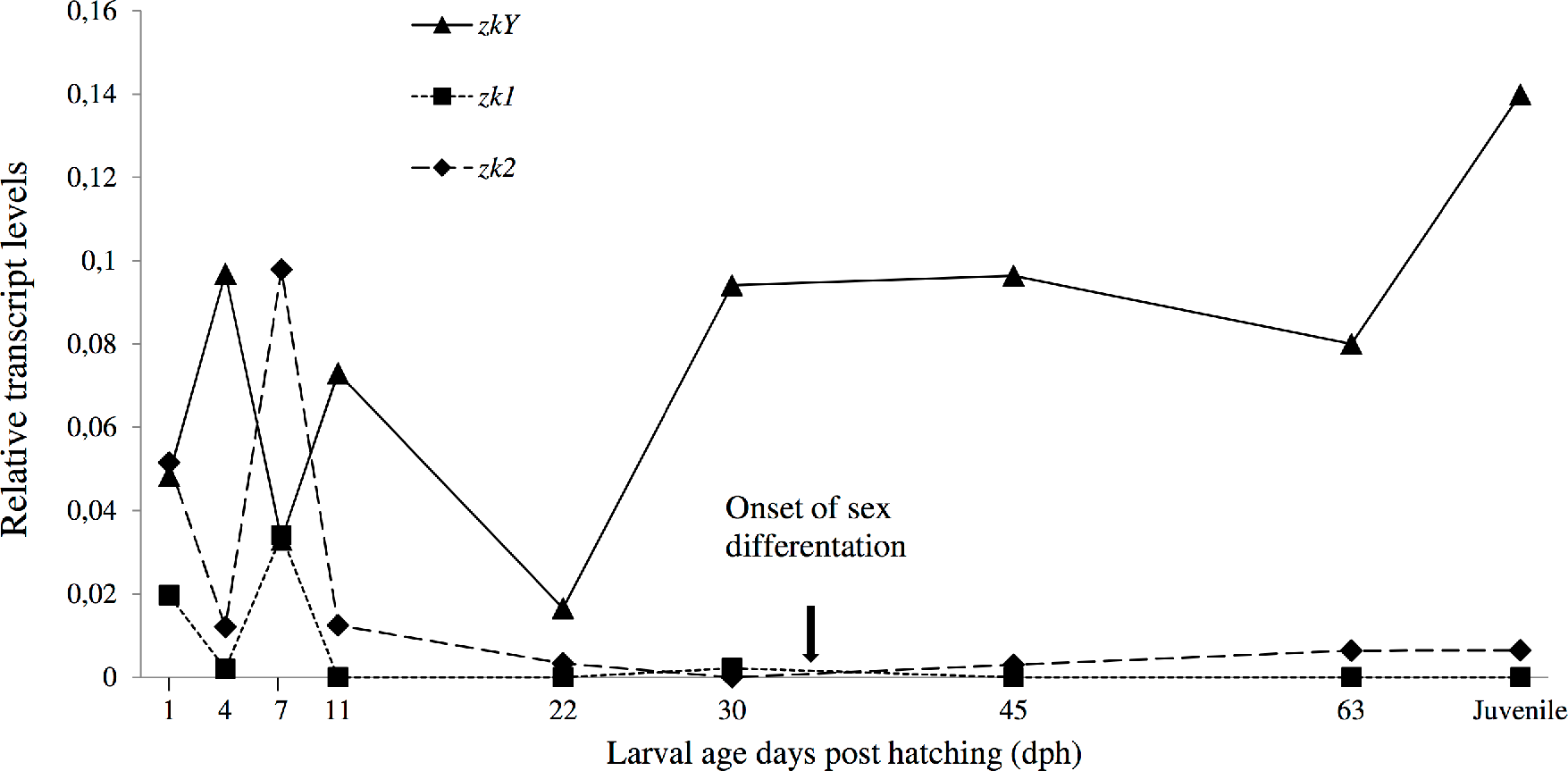
Larval expression of *zkY, zk1* and *zk2* from hatching to juvenile stage quantified as number of RNA reads. The commencement of sex differentiation is indicated [24].

The increased larval expression of the male-specific *zkY* gene prior to the onset of sex differentiation suggests that this novel zinc knuckle protein functions as the male determinant in Atlantic cod. Multiple germline-specific RNA regulatory proteins are essential for the germ cell specification, maintenance, migration and proliferation [57]. While this regulatory network seems to be evolutionary conserved in worms, flies and mammals, the key role played by different sex determinants in the control of male germ cell proliferation in fish suggests the involvement of lineage-specific regulators [6]. Intriguingly, knockdown of the germline zinc knuckle helicases *glh1-2* in *C. elegans* and the *vasa* homolog in mice resulted in male infertility [58-60]. The loss of zinc knuckles in the RNA-binding Vasa in vertebrates and insects was suggested to have coincided with the emergence of hitherto unknown zinc-knuckle cofactors conferring target specificity [61]. We, therefore, propose that cod zkY activates male sex determination in Atlantic cod by interacting with the specific RNA targets governing germ cell proliferation.

### X- and Y-sequences in other gadoid species

To elucidate the emergence of the sex determining region in other cod fishes we scanned draft genome assemblies from 12 species within the family *Gadidae* [62] for sequences matching the Atlantic cod Y- sequence. In its entirety, the 9,149 bp male-specific region showed only partial hits suggesting that the Y- sequence is absent or incorrectly assembled in these draft assemblies. This result highlights two important issues associated with using draft genome assemblies to characterize heteromorphic sex determining regions. First, analyses are hampered if metadata is incomplete and important basic information such as the gender of sequenced individuals is missing, as in the case of draft assemblies submitted by Malmstrøm et al. 2017 [62]. Secondly, assemblies constructed purely from short paired-end reads may be severely impacted by genome heterogeneity, repeats, and structural variation. Therefore, to find evidence for the sex determination region in other codfish species we were obliged to use a PCR based approach. Although the multi-species sequence alignments were highly fragmented, a close examination revealed isolated regions within and immediately outside the male-specific region of Atlantic cod, which were, to some extent conserved across species. These regions allowed us to develop primers, which amplified a specific fragment of the Y-sequence in both Greenland cod (*Gadus macrocephalus ogac*) and Arctic cod (*Arctogadus glacialis*), in addition to Atlantic cod (see alignment in S7 file). The X-sequence was efficiently amplified and sequenced in Greenland and Arctic cod, Polar cod (*Boreogadus saida*), haddock (*Melanogrammes aeglefinus*) and burbot (*Lota lota*) (see alignment in S8 file). This result documents that the Y-specific sequence was present prior to the separation of Arctic cod from Atlantic cod more than 7.5 MYA [62, 63] while the presence of X-sequence in burbot suggests that this sex-determining mechanism predates the *Lotinae* – *Gadinae* split 45 MYA.

A diagnostic test for distinguishing male and female Atlantic cod is potentially useful for researchers and the emerging aquaculture industry. To efficiently determine gender, we developed a simple PCR reaction including two different forward primers (one Y-sequence specific and the other matching a common sequence upstream of the X- and Y- regions) and a common reverse primer (annealing to a sequence downstream of the X- and Y-regions) resulting in a single band in females and double in males (Fig 4).

**Figure 4.**
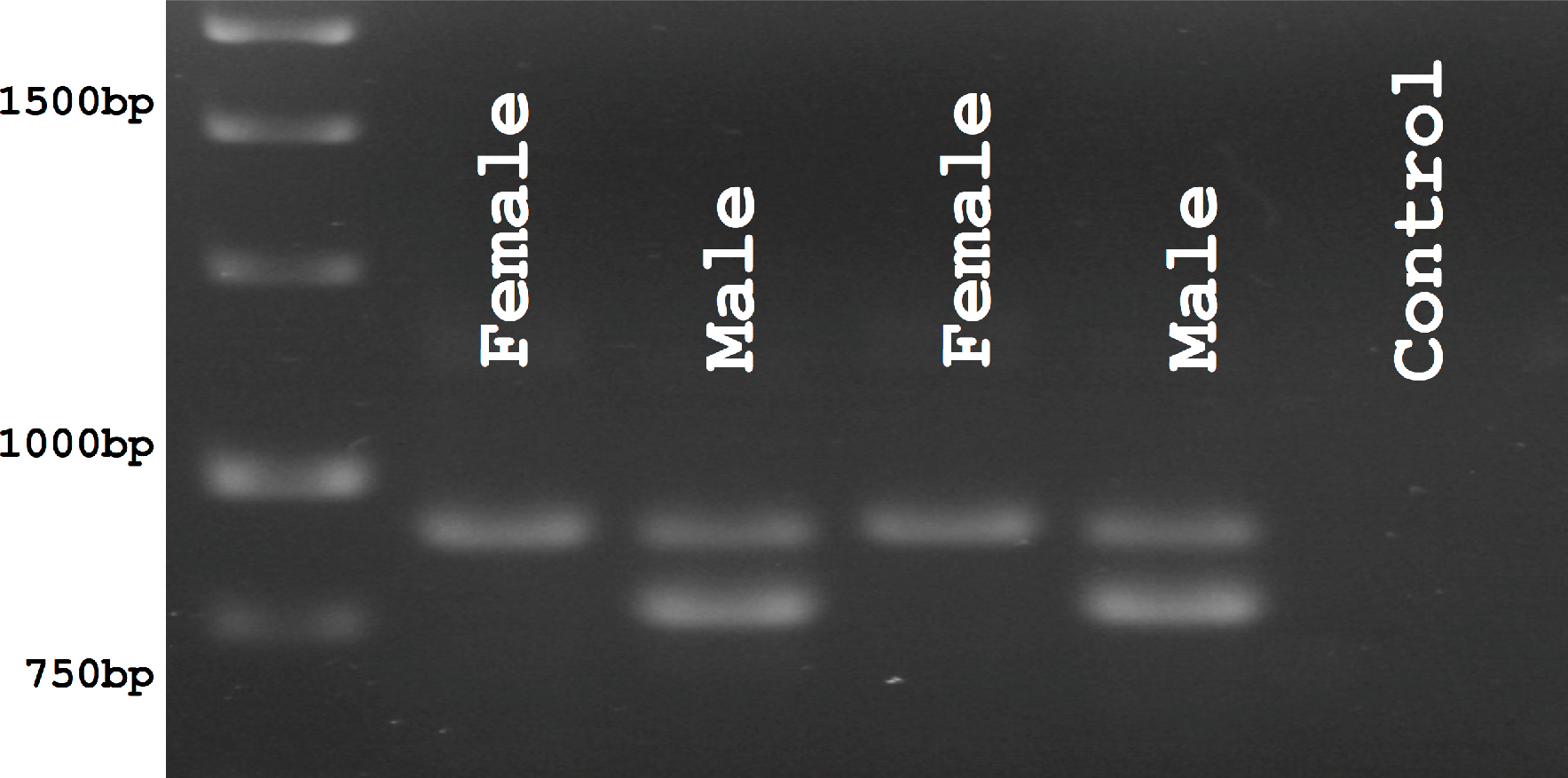
Agarose gel separation of PCR products from male and female Atlantic cod generated using sex-specific primer combinations.

Sex determination is a complex process involving the coordinated actions of multiple factors, and a greater understanding of what genes are involved in the sex determination cascade is needed. Furthermore, this understanding must be developed in multiple species to learn whether these processes are species or taxa specific and to improve our knowledge of the evolution of sex determination. The main focus of this paper has been to understand the mechanistic basis for sex determination in Atlantic cod. Our results strongly indicate that the single *zkY* gene present in the Y-sequence on LG11 functions as the major masculinization regulator in Atlantic cod and documents the presence of X- and Y- sequences in other gadoids. Our interpretation has been complicated by the presence of multiple, similar genes distributed across the genome and a collapse of X- and Y-sequences in the available reference genomes and the fact that *zkY* has not been recruited from the conserved genes within the sex differentiation cascade. Therefore, an emphasis must be placed on producing high quality reference genomes if our understanding of teleost sex determination is to advance.

## Materials and methods

### Sampling and Illumina whole-genome sequencing

Tissue was collected from 49 males and 53 females from the National Atlantic cod breeding program maintained by Nofima, Tromsø, Norway (available at https://www.ebi.ac.uk/ena, PRJEB12803 and S9 File). The fish were parents in year classes 2005 or 2006 and represented the second generation of cod produced in captivity. The original broodstock in the base population were sampled from different geographical areas along the Norwegian coast [64]. DNA was extracted using the DNeasy kit manufactured by QIAGEN (Germany) according to manufacturer’s instructions. Libraries were prepared using the Truseq Library prep kit (Illumina, San Diego, USA), and sequenced using an Illumina HiSeq 2500 instrument. A total of 13.1 billion paired-reads (2 x 100 PE) were produced, averaging 129 million paired reads per individual with coverage from 12.4 to 19.0X.

### Construction of gadMor2.1 assembly and variant detection

The gadMor2.1 assembly was constructed by integrating a dense linkage map with two public draft assemblies (NEWB454 and CA454ILM https://figshare.com/articles/Transcript_and_genome_assemblies_of_Atlantic_cod/3408247) generated from sequencing the same male. The assemblies were constructed using different combinations of short-read sequencing data and different assembly programs (Newbler and Celera respectively) and displayed different qualities with NEWB454 having longer scaffold N50s and more gaps than CA454ILM [36]. The linkage map containing 9354 SNP markers was produced after genotyping a pedigree of 2951 fishes. Chimeric scaffolds were broken at junctions between contigs containing SNPs from different LGs. Overlapping scaffolds were identified by comparing SNPs mapping to both assemblies and were merged using coordinates from alignment with LASTZ [65] generating scaffolds that were used to build the final chromosome sequences. Finally, all scaffolds were oriented, ordered and concatenated into a new chromosome sequence based on information from the linkage map.

Variants were detected by first filtering raw reads from each individual using Trimmomatic v0.32 [66], and subsequently aligning reads to the unmasked gadMor2.1 assembly using Bowtie2 v2.3.2 with the parameter ‘sensitive’ [67]. The resulting BAM files were merged by gender using Sambamba v0.6.5 [68] and GATK HaplotypeCaller v3.8 [69] was used to call variants with the following parameters: gt_mode DISCOVERY, minPruning 3. A Perl script was used to report SNPs, and their positions, that display heterozygous genotypes in all males and homozygous genotypes in all females.

### Read depth differences between males and females

For each sex, quality filtered reads from males (n=49) and females (n=53) were combined to generate two gender-specific bed files. Bedtools genomecov version 2.22.0 [70] was then used to count read depth at each position in the gadMor2.1 assembly. A Perl script was then used to calculate the average read depth per individual across successive 1000bp intervals with an overlap of 500bp (positions with 0 coverage in males and females were ignored). For each average, the absolute difference between the larger and smaller number was calculated and plotted. Across the genome, the region with the most extreme read depth differences was seen on LG11.

### Construction of Y- and X-sequences

A nested PCR strategy was used to amplify the Y-specific sequence lacking from the gadMor2.1 assembly. The initial reaction used the following primers and conditions, (Fwd; 5’-CCTAAACCACAGTCCTGGGC-3’, Rev; 5’-ACATTGTGCACACACATTGTATC-3’, PrimeSTAR GXL DNA Polymerase (Takara, Japan), cycling 30 times with 10s at 98°C, 15s at 57°C, 3 min at 68°C, final extension 72°C for 10 min. PCR-product was used as template in a subsequent reaction using the same conditions but with the nested PCR primers (Fwd; 5’-CTCTGTAGTTTGTGGTGGGGT-3’, Rev; 5’- ACATGACGAATGGCCTCCTTT-3’). The resulting PCR product was purified using a QIAquick PCR purification kit from QIAGEN (Germany) and quantified. DNA was prepared for sequencing using a Nextera XT kit from Illumina (USA) according to manufacturer’s instructions and the resulting library sequenced using 2 x 250 nt sequencing chemistry using a MiSeq (Illumina, USA) (S10 File). After quality trimming the resulting reads were anchored to the existing flanking Y-sequences using Gap filler [71]. The resulting sequence contained a single gap that was filled by Sanger sequencing a PCR product generated using the following PCR primers and conditions (Fwd; 5’-ACACAACGCAGAGTCTGTCC-3’, Rev: 5’- TCAGCTAGTCTCGCAATGGC-3’, denaturation 15 min at 95°C, then cycling 30 times with 10s at 98°C, 15s at 98°C, 60s at 68°C, final extension 10 min at 72°C). An X-sequence was constructed from Sanger sequencing a PCR product produced using primers annealing up- and down-stream of the Y-sequence.

### Generation X- and Y-sequences in other gadoids

Sequence alignments of Atlantic cod sex determining region with public assembly data from 12 gadid species [62], using MuMmer version 3.23 [72], identified four rather short regions that showed good conservation across species, including two regions within the Y-sequence (B; 12,520,541-12,520,838 and C; 12,527,782-12,5280,27) and two regions (A; 12,518,827-12,519,231 and D; 12,528,480-12,529,285) immediately flanking the X- and Y-sequences. Primers were designed using Primer3 [73] to amplify from within the Y-sequence conserved sequence C to flanking sequence D (i.e. a Y-specific PCR) and from flanking sequence A to D (an X-sequence specific PCR). Products from these reactions were Sanger sequenced and aligned using MAFFT v7.402 [74] (S7 file and S8 file).

### Diagnostic sex-test

A sex distinguishing duplex PCR was designed for Atlantic cod using the following primers and conditions (Fwd A; 5’-ACACACGGTCTGCTGTAGTG-3’, Fwd C; 5’- GGAGGGGAATTGTACAAACACG-3’, Rev D; 5’-GTGTGCCAAATGGATGCCAA-3’), denaturation for 15 min at 95°C, cycling 35 times with 30s at 94°C, 60s at 55°C, 60s at 72°C, final extension 72°C for 10 min.

### Nanopore Sequencing

High-molecular weight DNA was extracted using a phenol-chloroform method based on [75]. Sequencing libraries were prepared using SQK-RAD003 and SQK-LSK108 kits and protocols from Oxford Nanopore Technology (Oxford, UK) and sequenced on a MinION device generating a combined total of 3.9Gb sequence data with an N50 read length of 4.2kb. Reads were aligned to the gadMor2.1 assembly using GraphMap aligner v0.5.2 [76].

### Transcript profiling of cod larvae

Cod larvae were hatched at the National Atlantic Cod Breeding Centre in Tromsø, Norway, after the incubation of fertilized eggs in seawater rearing tanks at 4.5 °C. For 1 and 7 dph larvae, RNA was extracted from a pool of 10 individuals using the QIAGEN AllPrep DNA/RNA/miRNA Universal kit (QIAGEN; Germany). Samples were prepared for sequencing using TruSeq Stranded mRNA kit from Illumina (USA) and sequenced using an Illumina HiSeq 2000 to produce 362 million reads. For the samples 12 and 35 dph samples, RNA was extracted using an RNeasy kit (QIAGEN, Germany) prepared for sequencing using TruSeq Stranded mRNA kit from Illumina (USA) and sequenced using an Illumina MiSeq (2 x 250nt) to produce 4.4 million reads. All reads were trimmed using Trimmomatic v0.32 [66] before further analysis.

Total available short read data (http://www.ebi.ac.uk/ena, PRJEB18628; and https://www.ncbi.nlm.nih.gov/sra/?term=SRP056073) was binned based on days before hatching (dph) before being aligned to the gadMor2.1 assembly using star aligner STAR v2.3.1z12 as described previously. Potential transcripts were constructed using stringtie v1.3.3 [78], while cuffmerge v2.2.1 was used to produce a GTF file containing key metrics. FPKM values for zkY, zky1, zky2, were calculated using only those reads with a mapping quality of ≥30.

### Gene annotation

Data from various public sources was used to build gene models including (i) 3M transcriptome reads generated using GS-FLX 454 technology and hosted at NCBI’s SRA (https://www.ncbi.nlm.nih.gov/sra/?term=SRP013269), (ii) >250K ESTs hosted by NCBI (https://www.ncbi.nlm.nih.gov/nucest) (iii) 4.4M paired-end mRNA sequences from whole NEAC larvae at 12 and 35 dph (https://www.ebi.ac.uk/ena, PRJEB25591), (iv) Pacbio reads from (https://www.ebi.ac.uk/ena, PRJEB18628), (v) 362M Illumina reads from 1 and 7 dph (https://www.ebi.ac.uk/ena, PRJEB25591) and (vi) approximately 1.7B Illumina reads from 4 - 63 dph as well as juvenile samples (https://www.ncbi.nlm.nih.gov/sra/?term=SRP056073). To enable model building, short Illumina reads (< 250nt) were mapped to the gadMor2.1 assembly using STAR v2.3.1z12. Because Illumina reads from 4 dph - juvenile were generated using an unstranded library, the parameter ´outSAMstrandField intronMotif’ was used in alignment. Long reads from PacBio were mapped using STARlong v2.5.2a [77] while 454 transcriptome reads were mapped using gmap v 2014-07-28 [78] with ‘–no-chimeras’ parameter in addition to default parameters. Cufflinks v2.2.1 [79] was used to assemble the reads into transcript models for all alignments except for data from 4 dph - juvenile stage samples where stringtie v1.3.3 [80] was used. Transcript models were merged using cuffmerge v2.2.1 [79].

Gene models were tested by performing (i) open reading frame (ORF) prediction using TransDecoder [81] using both pfamA and pfamB databases for homology searches and a minimum length of 30 amino acids for ORFs without pfam support, and (ii) BLASTP analysis (evalue <1e-10) for all predicted proteins against zebrafish (*Danio rerio*) (v9.75) and three-spined stickleback (*Gasterosteus aculeatus*) (BROADS1.75) annotations from Ensembl. Only gene models with support from at least one of these homology searches were retained. Functional annotation of the predicted transcripts was done using blastx against the SwissProt database.

### Modeling the zinc knuckle domain

The three-dimensional structure of the three different variants of the zinc knuckle domain were built using a threading approach which combines three-tridimensional fold recognition by sequence alignment with template crystal structures and model structure refining. The query-template alignment was generated by HHPRED (https://toolkit.tuebingen.mpg.de) and then submitted to the program MODELLER [82] as implemented in Discovery Studio (Dassault Systèmes BIOVIA). Query coverage and e-value score were considered to define a suitable template structure. The structure of nucleocapsid protein NCp10 of retrovirus MoMuLV, which contains a single Cys-X2-Cys-X4-His-X4-Cys zinc knuckle domain bound to the oligonucleotide d(ACGCC) was selected as template (https://www.rcsb.org/structure/1a6b). Zinc ions and oligonucleotides were explicitly considered through molecular modeling steps. Fifty models, optimized by a short simulated annealing refinement protocol available in MODELLER, were generated and their consistency was evaluated on the basis of the probability density function violations provided by the program. Stereochemistry of selected models was checked using the program PROCHECK [83]. Visualization and manipulation of molecular images were performed with Discovery Studio (Dassault Systèmes BIOVIA).

## Data accessibility

All supporting information is available at https://figshare.com/s/f37196e26871266be664.

## Acknowledgements

Thanks to Kim Præbel at The Arctic University of Norway (UiT) for providing DNA samples from haddock, Arctic cod, Greenland cod, Polar cod and burbot. Thanks also to Nicola Barson for her valuable, critical review of the manuscript.

## Supporting information

**S1 File.** Atlantic cod gadMor2.1 genome assembly.

**S2 File.** The position and the flanking sequences of the nine sex-specific SNPs on LG11 that is heterozygous in all 49 males and homozygous in all 53 females of Atlantic cod.

**S3 File.** The X-sequence and the Y- sequence, including cod *zkY*, in the gadMor2.1 assembly.

**S4 File.** Oxford nanopore sequences of Atlantic cod that aligned to the X- and Y- sequences in the gadMor2.1 assembly.

**S5 Table.** The position of the eight autosomal gene copies of cod *zkY* in gadMor2.1 assembly.

**S6 File**. MAFFT alignments of cod zkY and the eight autosomal copies in Atlantic cod.

**S7 File**. MAFFT alignments of the Atlantic cod Y-sequence with other gadoids.

**S8 File**. MAFFT alignments of the Atlantic cod X-sequence with other gadoids.

**S9 File.** 102 samples ids, sex and ENA accession id.

**S10a and S10b File.** The long-range miseq illumina data.

## References

1. Graves JA. Sex chromosome specialization and degeneration in mammals. Cell. 2006;124(5):901–14.

2. Bellott DW, Skaletsky H, Pyntikova T, Mardis ER, Graves T, Kremitzki C, et al. Convergent evolution of chicken Z and human X chromosomes by expansion and gene acquisition. Nature. 2010;466(7306):612–U3.

3. Peichel CL. Convergence and divergence in sex-chromosome evolution. Nat Genet. 2017;49(3):321–2.

4. Devlin RH, Nagahama Y. Sex determination and sex differentiation in fish: an overview of genetic, physiological, and environmental influences. Aquaculture. 2002;208(3-4):191–364.

5. Volff JN, Nanda I, Schmid M, Schartl M. Governing sex determination in fish: Regulatory putsches and ephemeral dictators. Sex Dev. 2007;1(2):85–99.

6. Kikuchi K, Hamaguchi S. Novel sex-determining genes in fish and sex chromosome evolution. Dev Dyn. 2013;242(4):339–53.

7. Nanda I, Kondo M, Hornung U, Asakawa S, Winkler C, Shimizu A, et al. A duplicated copy of DMRT1 in the sex-determining region of the Y chromosome of the medaka, *Oryzias latipes*. PNAS. 2002;99(18):11778–83.

8. Matsuda M, Nagahama Y, Shinomiya A, Sato T, Matsuda C, Kobayashi T, et al. DMY is a Y-specific DM-domain gene required for male development in the medaka fish. Nature. 2002;417(6888):559–63.

9. Myosho T, Otake H, Masuyama H, Matsuda M, Kuroki Y, Fujiyama A, et al. Tracing the emergence of a novel sex-determining gene in medaka, *Oryzias luzonensis*. Genetics. 2012;191(1):163–70.

10. Takehana Y, Nagai N, Matsuda M, Tsuchiya K, Sakaizumi M. Geographic variation and diversity of the cytochrome b gene in Japanese wild populations of medaka, *Oryzias latipes*. Zoolog Sci. 2003;20(10):1279–91.

11. Kamiya T, Kai W, Tasumi S, Oka A, Matsunaga T, Mizuno N, et al. A trans-species missense SNP in Amhr2 is associated with sex determination in the tiger pufferfish, *Takifugu rubripes* (fugu). Plos Genet. 2012;8(7):e1002798.

12. Li M, Sun Y, Zhao J, Shi H, Zeng S, Ye K, et al. A tandem duplicate of anti-mullerian hormone with a missense SNP on the Y chromosome is essential for male sex determination in nile tilapia, *Oreochromis niloticus*. Plos Genet. 2015;11(11):e1005678.

13. Roberts NB, Juntti SA, Coyle KP, Dumont BL, Stanley MK, Ryan AQ, et al. Polygenic sex determination in the cichlid fish *Astatotilapia burtoni*. BMC Genomics. 2016;17(1):835.

14. Rondeau EB, Messmer AM, Sanderson DS, Jantzen SG, von Schalburg KR, Minkley DR, et al. Genomics of sablefish (*Anoplopoma fimbria*): expressed genes, mitochondrial phylogeny, linkage map and identification of a putative sex gene. BMC Genomics. 2013;14:452.

15. Nakamura Y, Yamamoto Y, Usui F, Mushika T, Ono T, Setioko AR, et al. Migration and proliferation of primordial germ cells in the early chicken embryo. Poult Sci. 2007;86(10):2182–93.

16. Zust B, Dixon KE. Events in germ-cell lineage after entry of primordial germ-cells into genital ridges in normal and uv-irradiated xenopus-laevis. J Embryol Exp Morph. 1977;41(Oct):33–46.

17. Nakamura M, Kobayashi T, Chang XT, Nagahama Y. Gonadal sex differentiation in teleost fish. J Exp Zool. 1998;281(5):362–72.

18. Kobayashi T, Matsuda M, Kajiura-Kobayashi H, Suzuki A, Saito N, Nakamoto M, et al. Two DM domain genes, DMY and DMRT1, involved in testicular differentiation and development in the medaka, *Oryzias latipes*. Dev Dyn. 2004;231(3):518–26.

19. Haugen T, Almeida FF, Andersson E, Bogerd J, Male R, Skaar KS, et al. Sex differentiation in Atlantic cod (*Gadus morhua* L.): morphological and gene expression studies. Reprod Biol Endocrin. 2012;10:47.

20. Saillant E, Chatain B, Menu B, Fauvel C, Vidal MO, Fostier A. Sexual differentiation and juvenile intersexuality in the European sea bass (*Dicentrarchus labrax*). J Zool. 2003;260:53–63.

21. Herpin A, Schindler D, Kraiss A, Hornung U, Winkler C, Schartl M. Inhibition of primordial germ cell proliferation by the medaka male determining gene Dmrt I bY. BMC Dev Biol. 2007;7:99.

22. Fujimoto T, Nishimura T, Goto-Kazeto R, Kawakami Y, Yamaha E, Arai K. Sexual dimorphism of gonadal structure and gene expression in germ cell-deficient loach, a teleost fish. PNAS. 2010;107(40):17211–6.

23. Wargelius A, Leininger S, Skaftnesmo KO, Kleppe L, Andersson E, Taranger GL, et al. Dnd knockout ablates germ cells and demonstrates germ cell independent sex differentiation in Atlantic salmon. Sci Rep-Uk. 2016;6:21284.

24. Goto R, Saito T, Takeda T, Fujimoto T, Takagi M, Arai K, et al. Germ cells are not the primary factor for sexual fate determination in goldfish. Dev Biol. 2012;370(1):98–109.

25. Cutting A, Chue J, Smith CA. Just how conserved is vertebrate sex determination? Dev Dyn. 2013;242(4):380–7.

26. Matsuda M, Sakaizumi M. Evolution of the sex-determining gene in the teleostean genus *Oryzias*. Gen Comp Endocr. 2016;239:80–8.

27. Herpin A, Schartl M. Plasticity of gene-regulatory networks controlling sex determination: of masters, slaves, usual suspects, newcomers, and usurpators. EMBO Rep. 2015;16(10):1260–74.

28. Yano A, Nicol B, Jouanno E, Quillet E, Fostier A, Guyomard R, et al. The sexually dimorphic on the Y-chromosome gene (sdY) is a conserved male-specific Y-chromosome sequence in many salmonids. Evol Appl. 2013;6(3):486–96.

29. Pandian TJ, Koteeswaran R. Ploidy induction and sex control in fish. Hydrobiologia. 1998;384:167–243.

30. Penman DJ, Piferrer F. Fish gonadogenesis. part I: Genetic and environmental mechanisms of sex determination. Rev Fish Sci. 2008;16:16–34.

31. Ottera H, Thorsen A, Peruzzi S, Dahle G, Hansen T, Karlsen O. Induction of meiotic gynogenesis in Atlantic cod, *Gadus morhua* (L.). J Appl Ichthyol. 2011;27(6):1298–302.

32. Whitehead JA, Benfey TJ, Martin-Robichaud DJ. Ovarian development and sex ratio of gynogenetic Atlantic cod (*Gadus morhua*). Aquaculture. 2012;324:174–81.

33. Ghigliotti L, Fevolden SE, Cheng CH, Babiak I, Dettai A, Pisano E. Karyotyping and cytogenetic mapping of Atlantic cod (*Gadus morhua* Linnaeus, 1758). Anim Genet. 2012;43(6):746–52.

34. Garcia-Souto D, Troncoso T, Perez M, Pasantes JJ. Molecular Cytogenetic Analysis of the European Hake *Merluccius merluccius* (Merlucciidae, Gadiformes): U1 and U2 snRNA Gene Clusters Map to the Same Location. Plos ONE. 2015;10(12):e0146150.

35. Star B, Torresen OK, Nederbragt AJ, Jakobsen KS, Pampoulie C, Jentoft S. Genomic characterization of the Atlantic cod sex-locus. Sci Rep-Uk. 2016;6:31235.

36. Torresen OK, Star B, Jentoft S, Reinar WB, Grove H, Miller JR, et al. An improved genome assembly uncovers prolific tandem repeats in Atlantic cod. BMC Genomics. 2017;18(1):95.

37. Thorvaldsdottir H, Robinson JT, Mesirov JP. Integrative Genomics Viewer (IGV): high-performance genomics data visualization and exploration. Brief Bioinform. 2013;14(2):178–92.

38. Summers MF, Henderson LE, Chance MR, Bess JW, Jr., South TL, Blake PR, et al. Nucleocapsid zinc fingers detected in retroviruses: EXAFS studies of intact viruses and the solution-state structure of the nucleocapsid protein from HIV-1. Protein Sci. 1992;1(5):563–74.

39. Williams MC, Gorelick RJ, Musier-Forsyth K. Specific zinc-finger architecture required for HIV-1 nucleocapsid protein’s nucleic acid chaperone function. PNAS. 2002;99(13):8614–9.

40. Guerrerio AL, Berg JM. Design of single-stranded nucleic acid binding peptides based on nucleocapsid CCHC-box zinc-binding domains. J Am Chem Soc. 2010;132(28):9638–43.

41. Benhalevy D, Gupta SK, Danan CH, Ghosal S, Sun HW, Kazemier HG, et al. The human CCHC-type zinc finger nucleic acid-binding protein binds G-rich elements in target mRNA coding sequences and promotes translation. Cell Rep. 2017;18(12):2979–90.

42. Mitra M, Wang W, Vo MN, Rouzina I, Barany G, Musier-Forsyth K. The N-terminal zinc finger and flanking basic domains represent the minimal region of the human immunodeficiency virus type-1 nucleocapsid protein for targeting chaperone function. Biochemistry. 2013;52(46):8226–36.

43. Michalek JL, Besold AN, Michel SLJ. Cysteine and histidine shuffling: mixing and matching cysteine and histidine residues in zinc finger proteins to afford different folds and function. Dalton T. 2011;40(47):12619–32.

44. Laity JH, Lee BM, Wright PE. Zinc finger proteins: new insights into structural and functional diversity. Curr Opin Struc Biol. 2001;11(1):39–46.

45. Zhu LY, Wilken J, Phillips NB, Narendra U, Chan G, Stratton SM, et al. Sexual dimorphism in diverse metazoans is regulated by a novel class of intertwined zinc fingers. Gene Dev. 2000;14(14):1750–64.

46. de Luis O, Lopez-Fernandez LA, del Mazo J. Tex27, a gene containing a zinc-finger domain, is up-regulated during the haploid stages of spermatogenesis. Exp Cell Res. 1999;249(2):320–6.

47. Ma K, Liao M, Liu F, Ye B, Sun F, Yue GH. Charactering the ZFAND3 gene mapped in the sex-determining locus in hybrid tilapia (*Oreochromis spp*.). Sci Rep-Uk. 2016;6:25471.

48. Bradley KM, Breyer JP, Melville DB, Broman KW, Knapik EW, Smith JR. An SNP-based linkage map for zebrafish reveals sex determination loci. G3-Genes Genom Genet. 2011;1(1):3–9.

49. Reichwald K, Petzold A, Koch P, Downie BR, Hartmann N, Pietsch S, et al. Insights into sex chromosome evolution and aging from the genome of a short-lived fish. Cell. 2015;163(6):1527–38.

50. Rondeau EB, Laurie CV, Johnson SC, Koop BF. A PCR assay detects a male-specific duplicated copy of anti-mullerian hormone (amh) in the lingcod (*Ophiodon elongatus*). BMC Res Notes. 2016;9:230.

51. Hattori RS, Murai Y, Oura M, Masuda S, Majhi SK, Sakamoto T, et al. A Y-linked anti-mullerian hormone duplication takes over a critical role in sex determination. PNAS. 2012;109(8):2955–9.

52. Meric C, Goff SP. Characterization of Moloney murine leukemia virus mutants with single-amino-acid substitutions in the Cys-His box of the nucleocapsid protein. J Virol. 1989;63(4):1558–68.

53. Urbaneja MA, McGrath CF, Kane BP, Henderson LE, Casas-Finet JR. Nucleic acid binding properties of the simian immunodeficiency virus nucleocapsid protein NCp8. J Biol Chem. 2000;275(14):10394–404.

54. Hashimoto H, Hara K, Hishiki A, Kawaguchi S, Shichijo N, Nakamura K, et al. Crystal structure of zinc-finger domain of Nanos and its functional implications. EMBO Rep. 2010;11(11):848–53.

55. Mark-Danieli M, Laham N, Kenan-Eichler M, Castiel A, Melamed D, Landau M, et al. Single point mutations in the zinc finger motifs of the human immunodeficiency virus type 1 nucleocapsid alter RNA binding specificities of the gag protein and enhance packaging and infectivity. J Virol. 2005;79(12):7756–67.

56. Johnsen H, Tveiten H, Torgersen JS, Andersen O. Divergent and sex-dimorphic expression of the paralogs of the Sox9-Amh-Cyp19a1 regulatory cascade in developing and adult atlantic cod (*Gadus morhua L*.). Mol Reprod Dev. 2013;80(5):358–70.

57. Cinalli RM, Rangan P, Lehmann R. Germ cells are forever. Cell. 2008;132(4):559–62.

58. Gruidl ME, Smith PA, Kuznicki KA, McCrone JS, Kirchner J, Roussell DL, et al. Multiple potential germ-line helicases are components of the germ-line-specific P granules of *Caenorhabditis elegans*. PNAS. 1996;93(24):13837–42.

59. Kuznicki KA, Smith PA, Leung-Chiu WMA, Estevez AO, Scott HC, Bennett KL. Combinatorial RNA interference indicates GLH-4 can compensate for GLH-1; these two P granule components are critical for fertility in *C-elegans*. Development. 2000;127(13):2907–16.

60. Tanaka SS, Toyooka Y, Akasu R, Katoh-Fukui Y, Nakahara Y, Suzuki R, et al. The mouse homolog of Drosophila Vasa is required for the development of male germ cells. Gene Dev. 2000;14(7):841–53.

61. Gustafson EA, Wessel GM. Vasa genes: emerging roles in the germ line and in multipotent cells. Bioessays. 2010;32(7):626–37.

62. Malmstrom M, Matschiner M, Torresen OK, Star B, Snipen LG, Hansen TF, et al. Evolution of the immune system influences speciation rates in teleost fishes. Nat Genet. 2016;48(10):1204–10.

63. Bakke I, Johansen SD. Molecular phylogenetics of gadidae and related gadiformes based on mitochondrial DNA sequences. Mar Biotechnol. 2005;7(1):61–9.

64. Bangera R, Odegard J, Nielsen HM, Gjoen HM, Mortensen A. Genetic analysis of vibriosis and viral nervous necrosis resistance in Atlantic cod (*Gadus morhua* L.) using a cure model. J Anim Sci. 2013;91(8):3574–82.

65. Harris RS. Improved pairwise alignment of genomic DNA: The Pennsylvania State University.; 2007.

66. Bolger AM, Lohse M, Usadel B. Trimmomatic: a flexible trimmer for Illumina sequence data. Bioinformatics. 2014;30(15):2114–20.

67. Langmead B, Salzberg SL. Fast gapped-read alignment with Bowtie 2. Nat Methods. 2012;9(4):357–9.

68. Tarasov A, Vilella AJ, Cuppen E, Nijman IJ, Prins P. Sambamba: fast processing of NGS alignment formats. Bioinformatics. 2015;31(12):2032–4.

69. McKenna A, Hanna M, Banks E, Sivachenko A, Cibulskis K, Kernytsky A, et al. The Genome Analysis Toolkit: a mapreduce framework for analyzing next-generation DNA sequencing data. Genome Res. 2010;20(9):1297–303.

70. Quinlan AR, Hall IM. BEDTools: a flexible suite of utilities for comparing genomic features. Bioinformatics. 2010;26(6):841–2.

71. Nadalin F, Vezzi F, Policriti A. GapFiller: a de novo assembly approach to fill the gap within paired reads. BMC Bioinformatics. 2012;13 Suppl 14:S8.

72. Kurtz S, Phillippy A, Delcher AL, Smoot M, Shumway M, Antonescu C, et al. Versatile and open software for comparing large genomes. Genome Biol. 2004;5(2).

73. Untergasser A, Cutcutache I, Koressaar T, Ye J, Faircloth BC, Remm M, et al. Primer3-new capabilities and interfaces. Nucleic Acids Res. 2012;40(15).

74. Katoh K, Rozewicki J, Yamada KD. MAFFT online service: multiple sequence alignment, interactive sequence choice and visualization. Brief Bioinform. 2017.

75. Russell JSaD. Molecular cloning: A laboratory manual third edition: Cold Spring Harbor Laboratory Press; 2001.

76. Sovic I, Sikic M, Wilm A, Fenlon SN, Chen S, Nagarajan N. Fast and sensitive mapping of nanopore sequencing reads with GraphMap. Nat Commun. 2016;7:11307.

77. Dobin A, Davis CA, Schlesinger F, Drenkow J, Zaleski C, Jha S, et al. STAR: ultrafast universal RNA-seq aligner. Bioinformatics. 2013;29(1):15–21.

78. Wu TD, Watanabe CK. GMAP: a genomic mapping and alignment program for mRNA and EST sequences. Bioinformatics. 2005;21(9):1859–75.

79. Trapnell C, Williams BA, Pertea G, Mortazavi A, Kwan G, van Baren MJ, et al. Transcript assembly and quantification by RNA-Seq reveals unannotated transcripts and isoform switching during cell differentiation. Nat Biotechnol. 2010;28(5):511–5.

80. Pertea M, Pertea GM, Antonescu CM, Chang TC, Mendell JT, Salzberg SL. StringTie enables improved reconstruction of a transcriptome from RNA-seq reads. Nat Biotechnol. 2015;33(3):290–5.

81. Haas BJ, Papanicolaou A, Yassour M, Grabherr M, Blood PD, Bowden J, et al. De novo transcript sequence reconstruction from RNA-seq using the Trinity platform for reference generation and analysis. Nat Protoc. 2013;8(8):1494–512.

82. Sali A, Blundell TL. Comparative protein modeling by satisfaction of spatial restraints. J Mol Biol. 1993;234(3):779–815.

83. Morris AL, MacArthur MW, Hutchinson EG, Thornton JM. Stereochemical quality of protein structure coordinates. Proteins. 1992;12(4):345–64.

